# Chloroplast DNA methylation in the kelp *Saccharina latissima* is determined by origin and influenced by cultivation

**DOI:** 10.1101/2022.12.02.518695

**Authors:** Lydia Scheschonk, Anne M.L. Nilsen, Kai Bischof, Alexander Jueterbock

**Affiliations:** University of Bremen, Marine Botany & MARUM, 28359 Bremen, Germany; Algal and Microbial Biotechnology division, Faculty of Biosciences and Aquaculture, Nord University, Universitetsalleen 11, 8026 Bodø, Norway

**Keywords:** Keywords: Chloroplast, organelle genome methylation, epigenetics, non-model organism, cultivation, aquaculture, marine macrophyte, marine algae

## Abstract

DNA cytosine methylation is an important epigenetic mechanism in genomic DNA, but absent in the chloroplast DNA of most land plants. We detected methylation in the chloroplast DNA of the kelp *Saccharina latissima*, a non-model macroalgal species of high ecological (wild populations) and economic importance. Since the functional role of the chloroplast methylome is yet largely unknown, we compared for the first time the chloroplast DNA cytosine methylation between wild and cultured kelp from different climatic origins (High-Arctic (79 °N) and temperate (54 °N), laboratory samples at 5 °C, 10 °C and 15 °C). Our results suggest genome-wide differences in methylated sites, and methylation level, between the origins, and a strong effect of cultivation. Functions related to photosynthesis showed differential methylation only between origins. Significant differences in methylation between cultivated and wild samples of genes related to transcription and translation were unique to the high-arctic samples. Both findings indicate that origin and cultivation strongly, but differently affect the chloroplast methylome. Similar methylomes for samples from the same origin – independent from whether they grow in the wild or in the lab – suggest that origin-specific methylation marks on the chloroplast genome are inherited. Given that DNA-methylation affects gene expression, our study suggests that lab-cultivation alters epigenetically determined kelp chloroplast characteristics at least to the same degree as ecotypic differentiation does, but likely on a much shorter timescale. This indicates the capacity for rapid non-genetic adaptation in the kelp *Saccharina latissima*.

## Introduction

Primary production, by which photoautotrophic organisms convert oxidised inorganic carbon into organic carbon via photosynthesis, is the one crucial process that ensures the existence of nearly all food webs, both terrestrial and marine. Since chloroplasts are presumed to have developed from a free-living cyano-bacteria to an endosymbiontic organelle (Nishimura 2010), they feature their own, albeit reduced, genome. Due to the reduced nature of their genome, the organelles are semi-autonomous, and depend on extra-organellar products for their internal processes (Wang et al. 2020). Epigenetic mechanisms include histone modification, non-coding RNA, and DNA methylation (Boquete et al. 2021). They have been shown to play vital roles in many cellular processes, such as regulation of gene expression, mRNA processing, timing of DNA replication, determination of chromatin structure, or silencing of transposons (Finnegan et al. 1998; Zhang et al. 2018). DNA cytosine methylation is the most stable epigenetic mechanism that can persist for multiple generations (Lämke and Bäurle 2017), thus is presumed to be most relevant for epigenetic adaptation. In contrast, histone modifications and ncRNAs are often only stable for several days or weeks (Lämke and Bäurle, 2017). However, the function of the chloroplast genome methylation is yet unclear, but intensely researched in the few species of mostly agricultural use that show chloroplast methylation, such as tomato and rice, or the unicellular green alga Chlamydomonas (Kobayashi et al. 1990; Nishimura et al. 1999; Muniandy et al. 2019). In marine primary producers, the first chloroplast genome methylation has recently been documented in the brown macroalga Saccharina japonica (Teng et al. 2021). Its functional relevance remains unknown.

Macroalgae are the predominant source of coastal marine primary production (Field et al. 1998; Filbee-Dexter 2020). Nearly exclusively restricted to the shorelines of the continents, macroalgae act as foundation species of marine rocky shore ecosystems, in addition to forming the base of food webs, and hence, have high and diverse ecological value (Bartsch et al. 2008; Teagle et al. 2017; Duffy et al. 2019). Their economic value is increasing exponentially as macroalgae cultivation gains momentum outside of Asia as an environmentally sustainable new blue economy with a multitude of applications (Cai et al. 2021). In 2019, marine macroalgae accounted for ∼30% of the global production of marine aquaculture commodities (120 million t), worth US$ 14.7 billion (Cai et al. 2021). However, in recent years macroalgae distributions have shifted northward in response to increasing (summer) water temperatures (Krause-Jensen et al. 2020; Bringloe et al. 2020). Heat waves affect both wild populations and aquaculture yields. In kelp, the respective thermal tolerance of gameto- and sporophyte stages seems to be the strongest factor defining their global distribution limits (Monteiro et al. 2019; Liesner et al. 2020). Temperature is a controlling factor in many biological processes such as photosynthetic carbon acquisition. Enzymatic activity of RubisCO, the carbon capturing enzyme in photosynthesis, increases by a factor of 7 between 5°C and 15°C (Cen and Sage 2005), and has been shown to be especially vulnerable to high temperatures. As the chloroplast genome carries several photosynthesis-relevant enzymes, investigating its methylation at different temperatures is particularly relevant when considering methylome- based temperature adaptation in accordance to the eco-genotype approach (Scheschonk et al. 2022). While epigenetic mechanisms have been extensively researched in terrestrial plants, knowledge about kelp epigenetics still is limited to a handful of studies (Cock et al. 2010; Fan et al. 2020a, b; Scheschonk et al. 2022), with a single study on chloroplast DNA methylation (Teng et al. 2021). In primary producers, the chloroplast genome has been presumed to be un-methylated, despite data suggesting otherwise (*Saccharina japonica*, Teng et al 2021; *Chlamydomonas reinhardtii*, Nishimura 2010;). Epigenetic mechanisms hold the key to rapid local adaptation, and hence the capacity for species survival and ecosystem stabilisation despite swift alterations in biotic and abiotic factors. Especially in the face of the staggering changes in local and global ecosystems due to the anthropogenic climate crisis, gaining knowledge in the field of eco-evolutionary dynamics is crucial. Our study is the first attempt to determine the influence of temperature, origin, and growth condition (wild *vs* cultivated) on the methylome of chloroplasts of the kelp *Saccharina latissima* (sugar kelp). In the northern hemisphere, this boreal-temperate marine brown alga, a congener species to *S. japonica* with a large latitudinal distribution range, is among the macroalgal species of both high ecological and economic value (Bartsch et al. 2008; Cai et al. 2021).

In this fundamental research study, we tested the following hypotheses:

1. *Saccharina latissima* features chloroplast genome methylations, as has been reported for *S. japonica* (Teng et al 2021).
2. The chloroplast methylome mirrors the nuclear methylome (Scheschonk et al. 2022) in showing epigenetic differentiation between origins that remains under cultivation, suggesting a trans-generational epigenetic memory.
3. The chloroplast methylome changes in response to temperature and lab cultivation, thus providing potential for rapid non-genetic adaptation to environmental change.

## Materials and Methods

Laboratory cultures, field samples, and ‘DNA- Extraction and MethlylRAD sequencing are described in more detail in Scheschonk et al (2022). Changes and specific adjustments to the present study were necessary during the bioinformatics and statistics part, and are detailed below (‘Bioinformatics and Statistics’).

### Laboratory cultures

As described in more detail in Scheschonk et al (2022), four cultures per origin (S1-S4, H1-H4; technical replicates) were obtained from mono-parentally fertilised clonal gametophyte cultures of *Saccharina latissima* (Linnaeus) C.E. Lane, C. Mayes, Druehl & G.W. Saunders from Helgoland, German Bight (Nordstrand, 54°11’18.9“N 7°54’14.1”E; HG1, HG2, HG3, HG4 in culture since 2014) and Spitsbergen, Svalbard (Hansneset, Kongsfjorden, 78°59’26.0“N 11°58’42.3”E; SG1, SG2, SG3, SG4 in culture since 2015; see table Suppl. Table 1) were kept at each of the three temperatures (5 °C, 10 °C, and 15 °C) under white light at 15 µmol m-2 s-1 (18 : 6 hours light : dark) from as early as initiation of fertilization. Only sporophytes with a fresh weight exceeding 120 mg (without stipe) were taken for DNA extraction.

### Field Samples

Adult sporophytes were sampled from Helgoland, German Bight (54°11’18.9“N 7°54’14.1”E), and from Spitsbergen, Svalbard (78°59’26.0“N 11°58’42.3”E) at the same locations from which the gametophytes for the laboratory grown sporophytes originated. From each sporophyte, 4 discs (ø=3 cm) were cut from the fronds, omitting the meristematic region, ‘midsection’, and reproductive tissues. Samples from lab and field were dried in cellulose bags stored in silica. In total 21 samples from the laboratory cultures, and 20 samples from the wild were analysed. In contrast to the field samples, our laboratory samples were uniparentally fertilised. However, given that chloroplast genes are transmitted solely from the maternal parent (matrilineally; Kuroiwa 1991, in Nishimura 2010), the chloroplast genome inheritance is identical in uniparentally and bi-parentally fertilised individuals. This circumstance enables a direct comparison between the chloroplast methylomes of the cultured and wild samples.

### Bioinformatics and Statistics of MethylRAD sequencing

#### Clean-up of raw data

DNA was extracted and MethylRAD sequencing libraries were prepared as described in detail in Scheschonk et al (2022). The sequences were de-multiplexed by sample and quality trimmed with TrimGalore! v 0.4.1 (https://www.bioinformatics.babraham.ac.uk/projects/trim_galore/). Bases with Phred-score <20 (--quality 20) were eliminated, the adapter sequence was removed (--stringency 3), and the terminal 2bp (both ends) were removed to eliminate any artefacts at the ligation position (--clip_R1 2 \ --three_prime_clip_R1 2). Reads were checked for fragment (over-) representation, base AT CG content bias, and read length with FastQC v0.11.8 before and after quality trimming.

#### In-silico digestion

For an overview of potential methylation sites, and as a backbone to map sequences to the reference chloroplast genome for the *S. latissima* chloroplast (Fan et al. 2020b) was digested *in silico* using the custom python script InSilicoTypeIIbDigestion_corrected.py (http://marinetics.org/2017/04/11/REdigestions.html) and settings for simulating the restriction enzyme FspEI. The *in silico* digestion found 270 recognition sites, where 69 were CCGG recognition sites, 61 CCTGG sites, 77 CCTGT sites, and 63 CCAGG sites.

#### Mapping and annotation

The sequenced reads were mapped to the *in silico* digested reference genome of the *S. latissima* chloroplast, combined with the *in silico* digested genome of the closely related species *Saccharina japonica* to account for reads belonging to the nuclear genome and avoid false mappings to the chloroplast genome for those reads. Reads were mapped using Burrows- Wheeler Aligner (BWA Version: 0.7.17-r1188) (Li and Durbin 2010). Duplicate mappings were excluded from further analyses by filtering out reads with a mapping quality <10 using samtools (v 1.9). A count table was created listing the number of reads that mapped back to each methylation site. For this, htseq-count (v 0.7.2) was called on each sample alignment with *in silico* digested fragment in gff3 format as count features. An additional table was made with counts normalized to ‘reads per million (RPM) = (Reads mapped per site ×10^6)/(Total number of mapped reads)’ to account for varying sequencing depths.

Only methylated sites that were covered in at least one library with coverage >2 were kept in count tables and RPM tables. The sites were annotated for their sequence context (CG or CHG), genomic region, gene ID numbers, and gene functions based on the gff3 annotation file for the *S. latissima* chloroplast genome from NCBI (Accession: MT151382.1) (Fan et al. 2020b).

#### Statistics

##### Principal Component Analysis

Differences in the chloroplast methylome were investigated with principal component analysis (PCA) on RPM tables using the R package ’FactomineR’ (Lê et al. 2008). Since lab and field samples separated clearly into distinct groups in the PCA, their methylomes were compared first for all samples, and then separated into origins (Helgoland and Spitsbergen). Within the lab and field samples, and for all samples, we compared origins (Helgoland and Spitsbergen). Within the lab samples, we compared between the treatment temperatures of 5°C, 10°C, and 15°C.

##### Outlier detection

The outlier function in the R package ’FactoInvestigate’ (Thuleau & Husson, 2020) detected two outliers, FH1 and FH9 (Helgoland field samples) in the PCA. The outliers showed few reads mapping to the genome, few methylated sites, and high RPM values for some of the methylated sites (Suppl. Fig. 1). Thus, FH1 and FH9 samples were excluded from all follow-up analyses. No outliers were found in the Spitsbergen samples. As both outliers belonged to the same group (Helgoland field samples), outlier removal led to a reduction in variance explained by the first two principal components from 69.9% to 28.9% (Suppl. Figs. 1-3).

##### Methylation levels

We tested for differences in DNA methylation levels as RPM and for differences in the numbers of methylated sites between populations, between treatment temperatures, and between lab and field samples, using Wilcoxon rank sum tests. P-values were corrected with the Benjamini-Hochberg method (Benjamini and Hochberg 1995) to control for the false discovery rate, and results with adjusted p values padj <0.05 were considered significant. The tests were performed using the R package ’rstatix’ v0.7.0 (Kassambara, 2021).

We tested for correlation between the numbers of quality trimmed reads, and the numbers of quality-trimmed reads that mapped back to the *Saccharina latissima* chloroplast genome (Suppl. Fig. 2C, 3C, 2D, 3D).

##### Differential methylation analysis

We tested for significant differential methylation with the R package ’DESeq2’ (Love et al. 2014) between ‘Helgoland’ *vs* ‘Spitsbergen’ in all samples (‘All’, or ‘both locations’), and in only field samples and only lab samples. Further, between ‘field’ *vs* ‘lab’ in all samples, and in only Helgoland samples, and only Spitsbergen samples. We also compared pairwise between temperature treatments: both origins combined (Helgoland + Spitsbergen 5 °C *vs* 10 °C; 5 °C *vs* 15 °C; 10 °C *vs* 15 °C), separately for Helgoland and Spitsbergen (5 °C *vs* 10 °C; 5 °C *vs* 15 °C; 10 °C *vs* 15 °C), and for each temperature (Helgoland vs Spitsbergen @ 5 °C; 10 °C; 15 °C). ’DESeq2’ normalised raw methylation counts before testing for differential methylation, and corrected p-values with the Benjamini-Hochberg method for controlling false discovery rates (Benjamini & Hochberg, 1995). Methylated sites with adjusted p-value padj < 0.05 were considered significantly differentially methylated.

##### GO term analysis

Gene ontology (GO) terms for biological processes, cellular components, and molecular functions were extracted from the database https://www.uniprot.org/ using Gene ID numbers (Suppl. Table 2-5). The R package ’topGO’ (Alexa & Rahnenführer, 2009) was used to test for enrichment of the GO terms.

##### Gene names and functions

Names and functions of differentially methylated genes were extracted from the chloroplast genome of *Saccharina latissima* published on NCBI (strain y-c14; https://www.ncbi.nlm.nih.gov/nuccore/MT151382.1?report=graph).

##### Analysis of epigenetic distance

Epigenetic distances among lab and field samples (cultivation) or among samples from Helgoland vs Spitsbeergen (origin) were estimated as euclidean distances among samples in the PCA . T. The Shapiro-Wilk indicated non-normal distribution for groups analysed for cultivation (p-value lab p < 0.0001, field p < 0.0001) and for origin (Helgoland p < 0.0001, Spitsbergen p < 0.0001). Hence, we tested for differences in epigenetic distances among the cultivation or origin categories using non-parametric Wilcox-rank-sum test (suppl. Table 6).

## Results

### Methylation levels and methylated sites

*In-silico* digestion identified 69 MethylRAD sites for CG methylation, and 201 for CHG methylation in the chloroplast genome of *Saccharina latissima*. 20.7% and 24.8% of these sites were methylated, respectively (Table 1).

**Table 1:** Potentially methylated sites for each sequence context, the mean number of methylated sites per sample and standard deviations for the means. Methylation level is the mean percentage of potential methylation sites methylated in samples, shown in the table with standard deviations (SD).

| Sequence Context | Potential Sites<br>(in-silico) | Methylated sites<br>(mean +/- SD) | Methylation level<br>(mean +/- SD) |
| --- | --- | --- | --- |
| CG | 69 | 14.3 ± 1.5 | 20.7% ± 2.2% |
| CHG | 201 | 49.8 ± 3.0 | 24.8% ± 1.5% |

**Table 2:** All conditions that were found to be differentially methylated by the DESeq2 analysis when analysing the different rearing temperatures of the laboratory samples. Most significances were detected between the origins at respective temperature (Helgoland vs Spitsbergen @ T5/10/15), but only 3 of those were identical to those detected when all lab samples were being analysed as a group (Fig. 3, Suppl. Table 8 (‘ST8 Temperature‘)). However, genes QKE47546.1 and QKE47431.1 (bold letters) showed significantly different methylation when analysing for temperature difference as factor (origin, T vs T).

| Comparison | Gene Product | Gene ID |
| --- | --- | --- |
| 5° vs 10° Helgoland | **acetolactate synthase large subunit** | **QKE47546.1** |
| 5° vs 15° Helgoland | **acetolactate synthase large subunit** | **QKE47546.1** |
| 5° vs 10° Spitsbergen | **photochlorophyllide reductase subunit B** | **QKE47431.1** |
| 5° Helgoland vs Spitsbergen | photosystem I P700 apoprotein A2 _(1) | QKE47459.1 |
|  | ATP synthase CF1 alpha chain | QKE47418.1 |
|  | acetolactate synthase large subunit | QKE47546.1 |
|  | photochlorophyllide reductase subunit B | QKE47431.1 |
| 10° Helgoland vs Spitsbergen | photosystem I P700 apoprotein A2 _(1) | QKE47459.1 |
|  | ATP synthase CF1 alpha chain | QKE47546.1 |
|  | photosystem I assembly protein | QKE47473.1 |
|  | photochlorophyllide reductase subunit B | QKE47431.1 |
| 15° Helgoland vs Spitsbergen | photosystem I P700 apoprotein A2 _(1) | QKE47459.1 |
|  | ATP synthase CF1 alpha chain | QKE47546.1 |

The number of methylated CG sites was significantly higher in Helgoland than in Spitsbergen samples both in laboratory samples (p < 0.001, Fig. 1 A) and in laboratory and field samples combined (p < 0.0001, Fig. 1 B), but no significant differences were found between treatment temperatures or between field and laboratory conditions for sites in CHG contexts (Suppl. Fig. 3 C, D).

**Figure 1:**
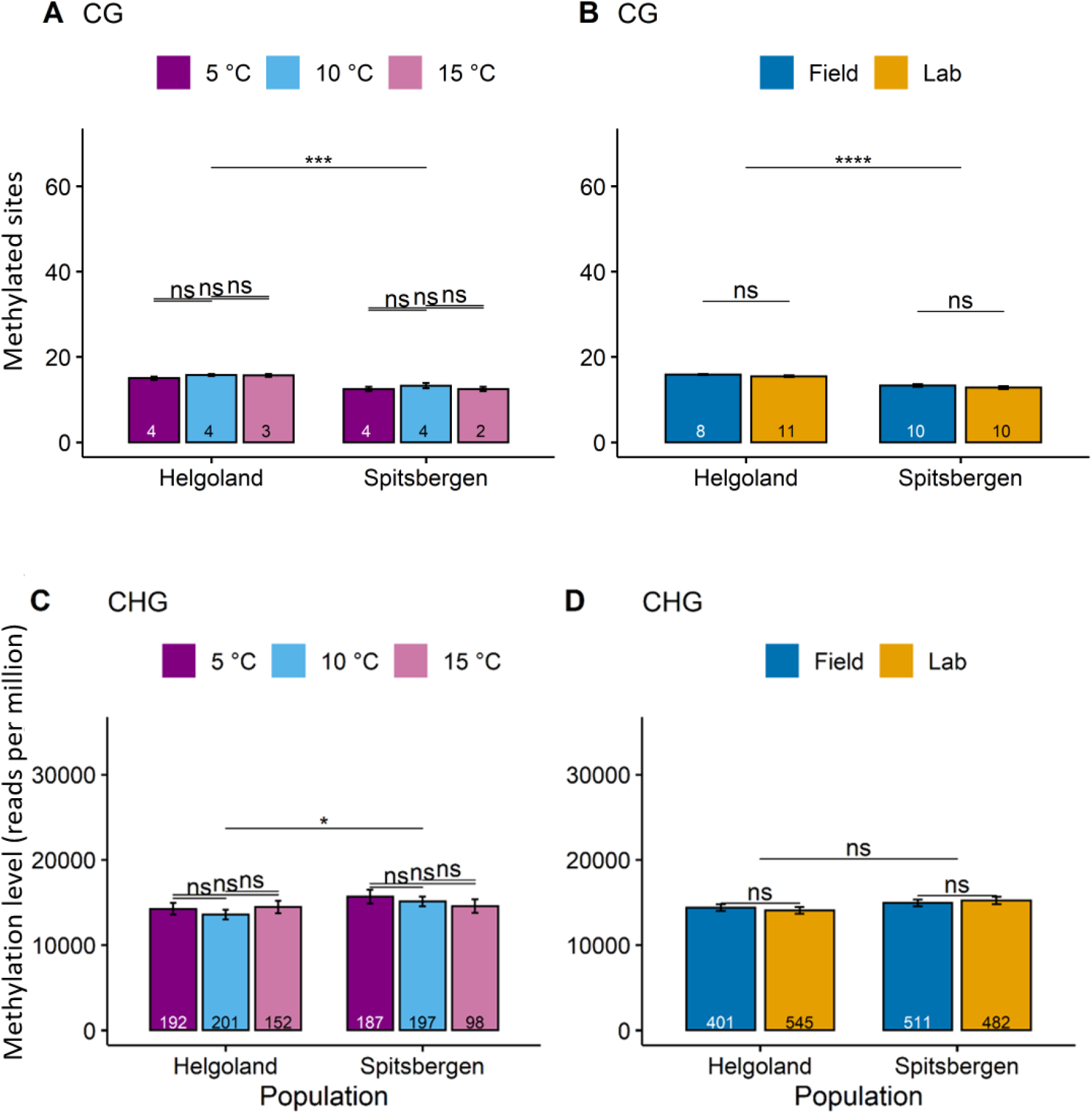
Significant differences in methylated sites and methylation level between conditions. Methylated CG sites (A, B), and methylation levels as reads per million (RPM) for CHG sites (C, D) between populations and temperatures in laboratory samples (A, C) and compared between lab and field samples (B, D). ns: not significant. Numbers at the bottom of the bars indicate sample size. Significance codes after p-value correction (Benjamini-Hochberg) for multiple comparisons: ‘< 0.05’: *, ‘< 0.01’: **, ‘< 0.001’: ***, ‘< 0.0001’: ****.

Methylation levels normalized to reads per million (RPM) showed no significant differences in CG contexts between any groups (Suppl. Fig. 4 A, B), but the Spitsbergen population showed higher methylation levels than the Helgoland population in the laboratory samples (p < 0.05, Fig. 1 C). There were no significant differences between field and lab conditions in CHG context (Fig. 1 D).

### Principal Component Analysis

Laboratory and field samples formed distinct groups along the first principal component with 95% confidence ellipses not overlapping when both origins were combined (Fig. 2 A). The first two principal components equally contributed about 14 % of the total sample variation, which was similar between the comparisons of ‘lab vs field’ (Fig. 2 A) and ‘Helgoland vs Spitsbergen’ (Fig. 2 B). However, for the origins (Fig. 2 B), the populations of Helgoland and Spitsbergen formed distinct groups along the second principal component when lab and field samples were combined (Fig. 2 B), while when field and laboratory samples were separately analysed for origin, Helgoland and Spitsbergen samples again split along the first component (Fig. 2 C, D). Temperatures in the laboratory samples (5°C, 10°C, 15°C) did not separate into distinct groups (Fig. 2 E).

**Figure 2:**
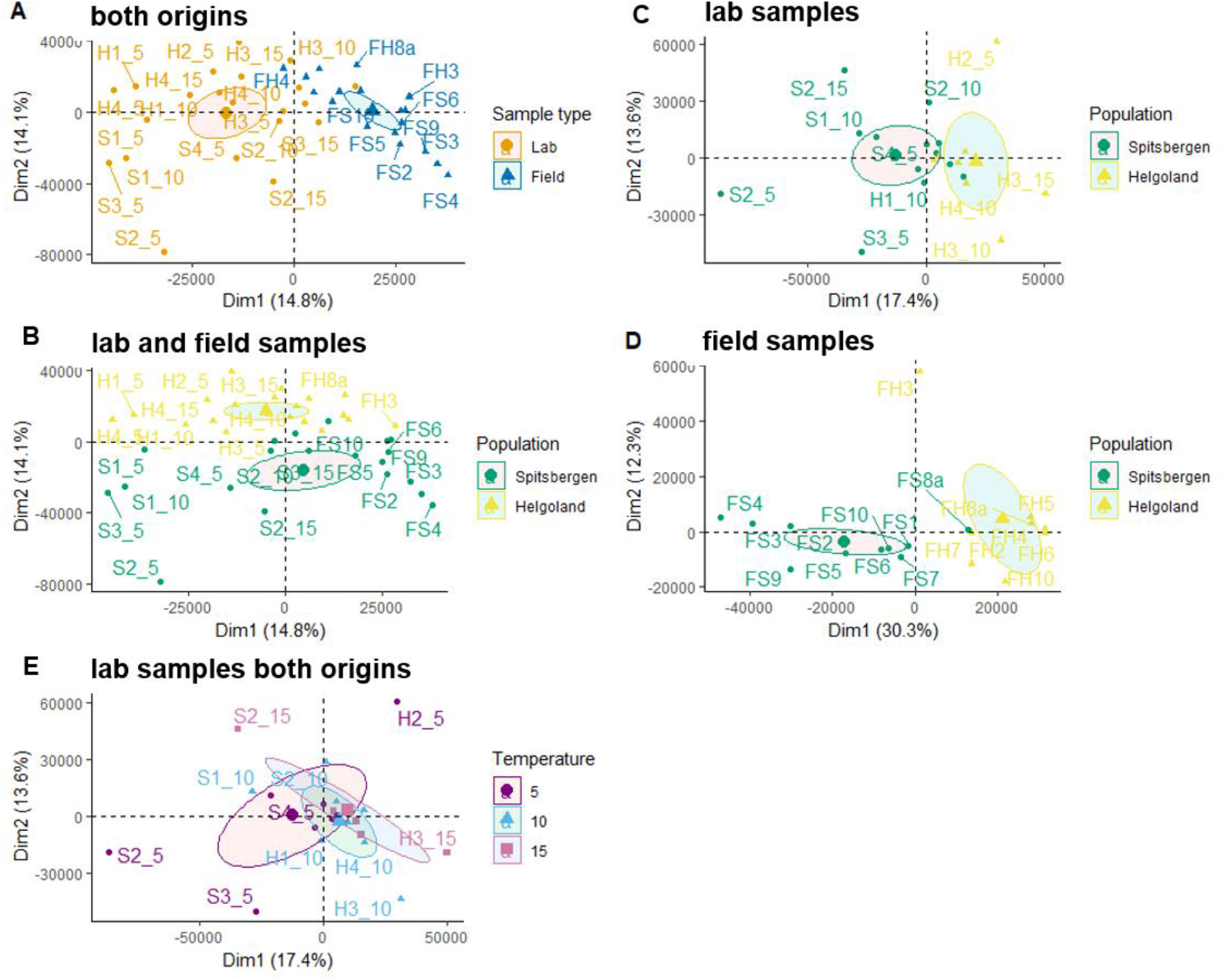
Principal Component Analysis for A: all samples, grouped by laboratory or field samples. B: laboratory samples grouped by origin. C: all samples, grouped by origin. D: field samples grouped by origin. E: laboratory samples of both origins grouped by temperature treatment. Ellipses represent 95% confidence intervals around the group medians.

### Differentially methylated sites and GO terms

Comparing the origins (Helgoland vs Spitsbergen, Fig. 3), differentially methylated sites were as often higher methylated in Spitsbergen samples as in Helgoland samples (Suppl.Tab.1-3). Of the eight genes with significant methylation differences between the origins (Fig. 3), only three genes (psaB_(1), atpA, yfc3) were identical to the genes with significant methylation differences between the origins at a respective temperature within laboratory samples (Table 2, see colour coding suppl. Table 7+8, sheets ‘ST7 Origin’ and ‘ST8 Temperature’).

**Figure 3:**
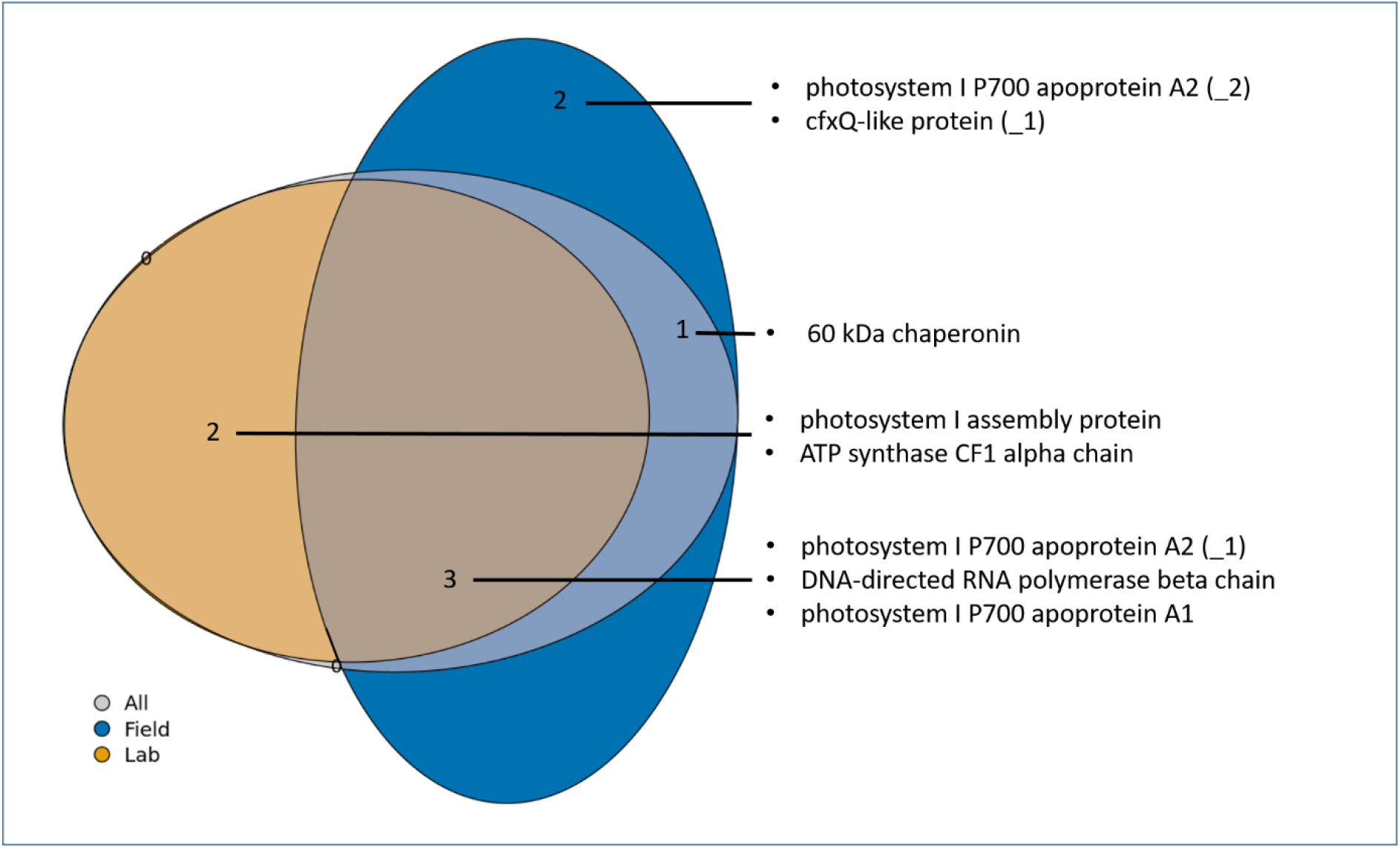
VennEuler diagram showing the numbers of differentially methylated sites between Helgoland and Spitsbergen when analysing field samples (blue), lab samples (orange), and all (lab+field, grey) samples (Suppl. Table 2-4). The labels give the product of the genes that were differentially methylated in the respective partitions. (_1) and (_2) depict a gene that showed differential methylation at two different sites. For the exact site position in the respective gene, see suppl. Table 7 (‘ST7 Origins’).

Of the 69 genes that were analysed comparing growth conditions (Fig. 4), 16 genes were differentially methylated between laboratory and field samples, two of which were also differentially methylated between origins (Fig. 3, Table 2). Many of those genes were differentially methylated at two different positions (suppl. Table 9, sheet ‘ST9 Cultivation’).

**Figure 4:**
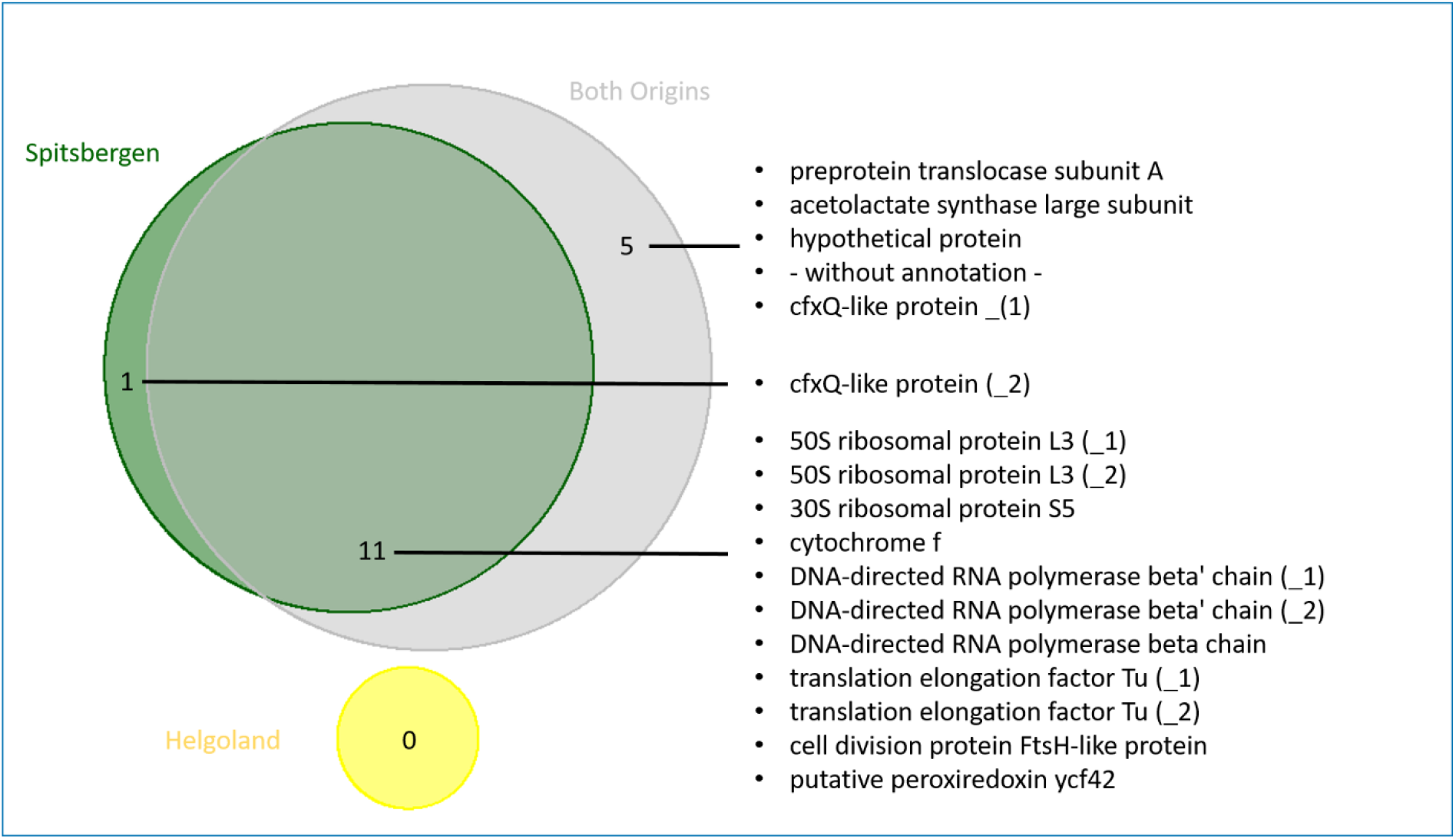
VennEuler diagram showing the numbers of differentially methylated sites between lab and field samples when analysing Helgoland samples (yellow), Spitsbergen samples (green), and the samples from both origins together (grey, Suppl. Table 5). The labels give the product of the genes that were differentially methylated in the respective partitions, with one gene without annotation record at the ncbi database. (_1) and (_2) refer to a gene that showed differential methylation at two different positions. For the exact position in the respective gene, see suppl. Table 9 (‘ST9 Cultivation’).

Two genes were differentially methylated between temperature treatments (Table 2), with ‘acetolactate synthase large subunit’ (QKE47546.1) at two comparisons: 5°C to 10 °C (padj < 0.0001), and 5 °C to 15°C (padj < 0.0017). All other significant differentially methylated genes were found between origins at different temperatures, but only partially overlapped with those obtained when analysing ‘all samples’ (Fig. 3, see colour codes in suppl. Tables 7 + 8 (sheets ‘ST7 Origins’ + ‘ST8 Temperature’)).

None of the extracted GOterms were significantly enriched according to the ’topGO’ enrichment analysis.

### Analysis of Epigenetic Distance

In both comparisons (‘cultivation’, ‘origin’), the epigenetic distances between the samples within respective groups were significantly different between groups (Fig. 5). For ‘cultivation’, distances were significantly higher in laboratory samples than in field samples (p < 0.0001, median field 153, lab 210), while for ‘origin’ the samples obtained from Spitsbergen showed significantly higher epi-distances than Helgoland (p < 0.0001, median Helgoland 171, Spitsbergen 190; Suppl. Table 6).

**Fig. 5:**
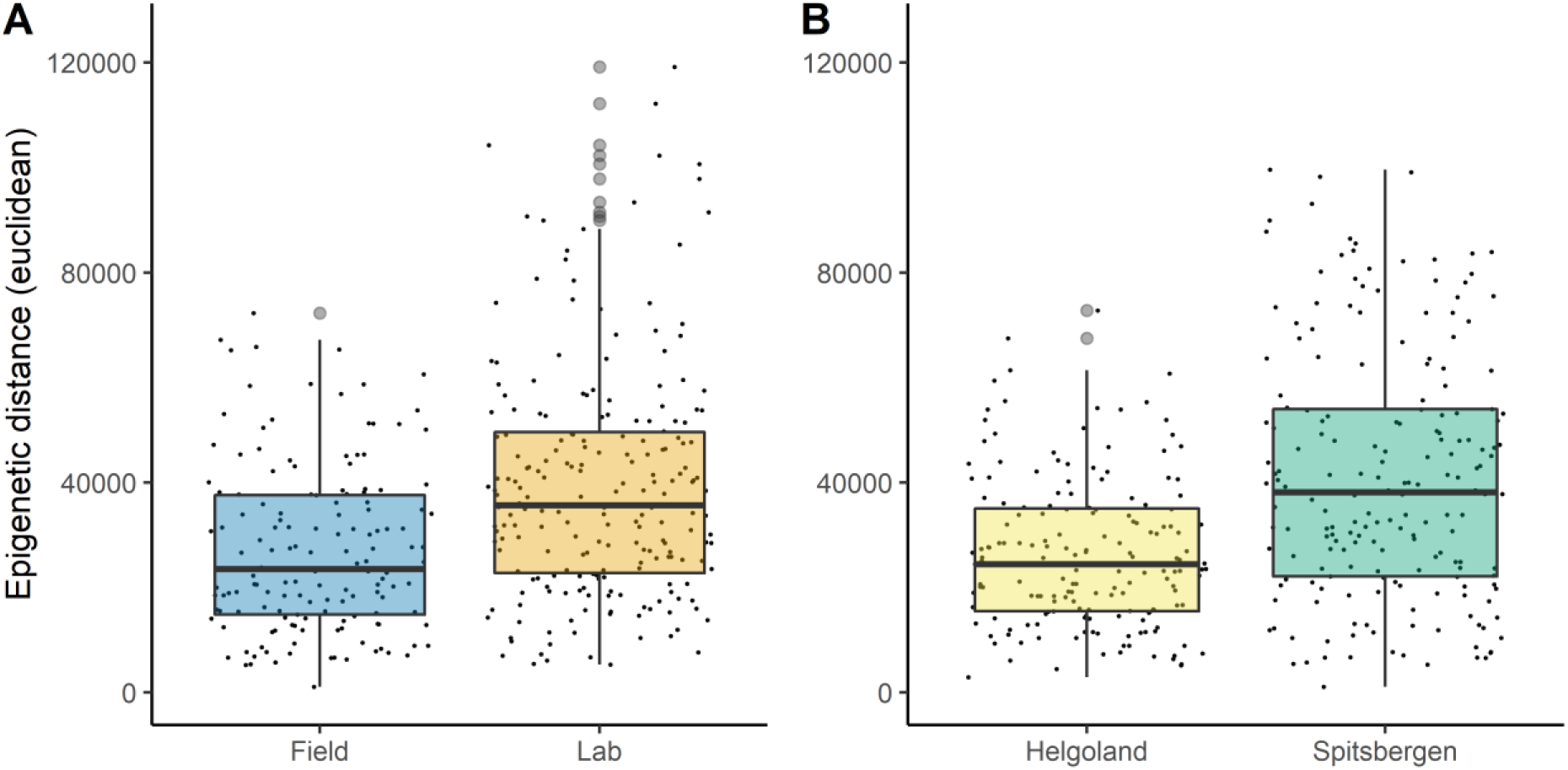
Epigenetic distance (Euclidean distance) for ‘cultivation’, Laboratory (orange) and Field (blue) samples (A), and for ‘origin’, Helgoland (yellow) and Spitsbergen (green) samples (B). The small black dots depict the pairwise distances between the individual samples of each group (from the PCA, Fig. 2 A,B). The light grey dots are outliers detected by the boxplot script, but were included during epi- distance analyses. The boxes represent the interquartile range (IQR) of the respective group, and the horizontal line within the boxes gives the median Euclidean distance.

## Discussion

Regarding Hypothesis 1, we could confirm a methylome in the chloroplast DNA of *Saccharina latissima* (Table 1). As with the nuclear genome (Scheschonk et al. 2022) the chloroplast methylome significantly differed between origins (Fig. 3, Table 2, Fig. 5 B), and growth conditions (Fig. 4, Fig. 5 A), suggesting a trans-generational epigenetic memory (Hypothesis 2). However, growth temperature had a smaller impact than expected (Table 2; Hypothesis 3). Differentially methylated sites between origins (Spitsbergen or Helgoland) were located in genes with functions related to photosynthesis or ATP synthesis. However, differentially methylated sites between growth conditions (laboratory or field) were predominantly located in genes related to translation and transcription (suppl. Table 7 + 9, sheets ’ST7 Origin’ + ‘ST9 Cultivation’; Fig. 3, Fig. 4). This suggests that the chloroplast methylome is origin-specific, indicating an epigenetic memory, while showing capacity of rapid adaptation to shifts in growth conditions via the epigenetic mechanism of cytosine methylation.

### Sample origin

Population origin had a strong impact on the methylome according to PCA clusters (Fig. 2 B, C, D), which was reflected in methylation levels, as well as in the number of methylated sites that significantly differed between origins (Fig 1 A, C). Photosystem I (PSI) was the component most affected by differences between the origins, as genes coding for varying aspects crucial to PSI significantly differed at a total of four sites in three genes (Fig. 3, Table 2), with the strongest effect on PSI P700 chlorophyll a apoprotein A2. The same gene was one of two genes differentially methylated between the origins in all analysed temperature conditions of the laboratory samples (Table 2, suppl. Table 8 sheet ‘ST8 Temperature’). Therefore, the difference can be assumed to be solely accounted for by origin. Part of the methylome was observed to be origin-specific even after years of gametophyte lab-cultivation. Methylation levels were significantly different between the origins (Fig. 1), independent from the rearing condition (laboratory or wild). As most of the differentially methylated genes between origins overlapped in field and lab samples (Fig. 3), this indicates the prevalence of a transgenerational epigenetic memory, which is further backed by PCA results. PCA in addition indicated the influence of cultivation on transgenerational memory, as origin played a more pronounced role when analysing laboratory and field samples separately (Fig. 2 C, D) *vs* combined (Fig. 2 B),. The difference was less pronounced in the laboratory than the field samples (Fig. 2 C *vs* D). The differentially methylated genes were exclusively related to functions of photosynthesis (Fig. 3), which likely can be explained by the latitudinal differences in the light regimes between origins. Light conditions in the High-Arctic are strongly seasonal, with absence of photosynthetically active radiation (PAR) for at least three months during the arctic winter (polar night), and continuous PAR during the arctic summer (polar day). In boreal-temperate regions, PAR is available throughout the year. Whether or not the differences observed here are genetically induced cannot be assessed by our setup, but the prevalence of the significant difference, detectable in the epigenetic signature that prevailed between origins through cultivation, is a strong indicator for the eco-genotype approach (Scheschonk et al 2022). This presumes that adaptation to a specific location is not solely a matter of genetic nature (fixed only in the DNA), but that epigenetic mechanisms, especially inheritable ones such as cytosine methylation, play an important role in local adaptation and eco-evolutionary dynamics of a species (Richards and Pigliucci 2020; Jablonka and Lamb 2015) that are partly independent from underlying genetic variation. Considering methylation and gene expression were shown to be negatively correlated in the *S. japonica* nuclear and chloroplast genes (Fan et al. 2020a; Teng et al. 2021) the (double-) methylation of the gene coding for apoprotein A2 (Fig. 3, suppl. Table 7 sheet ‘ST7 Origin’), for example, which is one of the crucial components of PSI, can be presumed to have a direct impact on the photosynthetic capacity which, according to our results, is likely to differ between origins. Like with the nuclear genome (Scheschonk et al 2022) this has crucial implications for cultivation processes as well as restoration efforts, as it shows the importance of suitable origin when considering local growth condition.

### Cultivation effects

PCA showed that cultivation affected the methylome to the same degree as origin (2 A, C). While lab and field samples showed no significant differences in genome-wide methylation level and number of methylated sites (Fig. 1), they displayed significantly different methylation levels for 23.19 % of the methylation sites (Fig. 4, suppl. Table 9 (sheet ‘ST9 Cultivation’). Most (12 out of 16) of the sites differentially methylated between rearing conditions had higher methylation levels in the lab than in field samples (Suppl. Table 2). This suggests those genes were higher expressed in the field samples (Fan et al. 2020a; Teng et al. 2021), possibly due to a higher variety of genes required in the more variable field conditions. Unexpectedly, epigenetic distance was higher in lab samples, even though they experience more uniform environmental conditions than field samples. When translated to transcription, a higher epigenetic distance is likely to result in more variable phenotypes. This might suggest that the kelp methylome diversifies rather under stable lab conditions than under variable field conditions, at least when the given cultivation conditions were not previously encountered. Potential explanatory factors cover the whole process of cultivation, from obtaining the sori, and securing the zoospores, or handling during the fertilisation process, to culture conditions at any stage.

However, origin as factor had the strongest influence on the differences in cultivated samples at gene level, while lab-field differences in methylation could only be detected in the Spitsbergen population (Fig. 4, Table 2). A factor that might as well explain differences in the methylomes between lab cultivated and wild kelp is the age of the sporophytes. The field sporophytes were adult, while the lab samples were juveniles with a size of approximately 12- 15 cm. In plants, cytosine methylation levels tend to increase with age, but these differences are most pronounced in meristematic areas (Dubrovina and Kiselev 2016), which were avoided in the kelp samples. Because meristematic regions were omitted, and older individuals (field samples) did not show higher methylation levels in the chloroplast genome, the likelihood that the results were affected by sporophyte age differences is low. Temperature as factor could be shown to have an origin-specific impact on the methylome during cultivation (Table 2; see next paragraph). However, which step, or factor, or combination thereof, during the cultivation process is likely to have caused the increase in epigenetic distance will have to be assessed in future studies.

Whether or not genotypic differences can result in the epigenetic differences that we observed has yet to be determined. In plants and algae, DNA cytosine methylation can be present in CG, CHG, and CHH sequence contexts, where H is any of the bases A, C, or T (Richards and Pigliucci 2020). Cytosine methylation can be heritable across several generations (Lämke and Bäurle 2017; Anastasiadi et al 2021), hence is a likely candidate for rapid non-genetic but heritable local adaptation. The inheritance of epigenetic traits induced by environmental factors has been repeatedly reported (Foust et al. 2016; Gugger et al. 2016; Keller et al. 2016). The difference in methylation between origins that persisted under laboratory conditions indicates epigenetic inheritance of a molecular memory of the environment. It suggests that methylation differences can remain stable across generations in cultivation, indicating that a transgenerational epigenetic memory likely is present in the *S. latissima* chloroplast genome.

### Temperature and the methylome

The temperature treatments did not form distinct clusters in the PCA. Unlike the nuclear methylome, the cultivated *S. latissima*s’ chloroplast methylome did not differ in methylation levels and numbers of methylated sites in response to the rearing temperatures of 5 °C, 10 °C and 15 °C. Significant differential methylation solely caused by temperature were detected in 2.9 % of the genes (Table 2). However, when analysing the origins at temperature level (Table 2), most differences between origins found at discrete temperatures were in other genes than those detected for the origin analysis. This again suggests the inheritance of an epigenetic memory in accordance with the eco-genotype approach (Scheschonk et al. 2022), as the methylomes reacted origin-specific at discrete temperatures. However, it cannot be ruled out whether these epigenetic differences have a genotypic base.

### Methylation levels compared with the nuclear genome

The methylation levels of sites in the chloroplast genome were ca. 100 times higher than levels in the nuclear genome as documented by Scheschonk et al. (2022). In the nuclear genome, the mean methylation level was 0.21% ± 0.13% SD (Scheschonk et al. 2022), while in the chloroplast, the levels were 20.7% ± 2.2% for CH contexts and 24.8% ± 1.5% for CHG contexts (Table 1). The higher methylation level in the chloroplast can be explained by the low numbers of potential methylation sites in the chloroplast genome (Table 1). The pattern of higher methylation in the chloroplast than the nuclear genome is repeated in both *S. japonica* and *S. latissima*, with some dissimilarities that could be caused by different sequencing methods and the inclusion of gametophytes in the studies of *S. japonica* (Fan et al. 2020a; Teng et al. 2021).

Photosynthesis is highly sensitive to temperature (Cen and Sage 2005). However, the main temperature response, in terms of methylation, is observable in the nuclear, but not in the chloroplast genome. The lack of statistically significant differences in the chloroplast genome as compared with the nuclear genome might partly be explained by the smaller number of genes. The chloroplast genome has a size of 130 kb and contains 139 protein-coding genes, three rRNA genes and 29 tRNA genes (Fan et al. 2020b), while the nuclear genome has a size of 545 Mb and contains 18,733 protein-coding genes (Ye et al. 2015). However, no significant differences were observed in neither methylation levels nor methylated sites between temperature treatments, despite the ∼ 100 x higher methylation level of the chloroplast. In contrast, in the nuclear genome, a response in methylation levels and numbers of methylated sites to warmer temperature treatments was observed, especially in the Helgoland samples (Scheschonk et al. 2022). Possible the lack of statistical significance indicates locations in the methylome where the methylation is highly conserved, and hence crucial on a functional level regardless of origin or external (new) impact. Aother possible explanation for the overall low amount of (differentially) methylated sites might be that the temperature exposure of the laboratory samples was not stressful for the physiological processes regulated by the chloroplast genome. The difference between nuclear and chloroplast methylome suggests that between 5 °C and 15 °C the modulation of temperature responses is solely regulated in the nuclear genome. In *Saccharina latissima*, it has been shown that temperature treatments affect the nuclear methylome, and that differences in the methylome exist between populations from different latitudes, suggesting that epigenetic modifications contribute to local adaptation (Scheschonk et al. 2022). In addition, significant differences in the nuclear methylome were found between wild and cultivated individuals (Scheschonk et al. 2022). Earlier research on *S. latissima* suggests that short-term acclimation to different growth temperatures can lead to changes in heat tolerance, and affect photosynthetic performance (Davison et al. 1991), which supports the temperature priming hypothesis suggested for the nuclear methylome. However, most of the differentially methylated genes were found between origins at discrete temperatures (5/10/15 °C), but were not detected when comparing the origins over all lab samples.

In the nuclear genome, most of the differentially methylated sites between lab and field samples and between populations were in non-genic repetitive contexts (transposable elements). In contrast, in the chloroplast genome, the differentially methylated sites were in coding regions of genes, except for very few sites in a promoter region (Suppl. Table 2-5). This is not surprising, given that chloroplast genomes are more condensed and have far fewer noncoding regions than nuclear DNA (Clegg et al. 1994).genomes.

## Conclusion

The chloroplast genome mainly contributes to processes of photosynthesis. Our results showed that sample origin and cultivation can strongly influence epigenetic mechanisms in the chloroplast, while temperature had a minor effect. Eco-phenotypes are likely developed in the chloroplast, as methylation differences in genes are prone to trigger differences in gene expression (Richards and Pigliucci 2020, Anastasiadi et al 2021). The scale-up in cultivation of *Saccharina latissima* during the last decade is likely to continue, and cultivation sites overall will likely shift northward, in accordance to the preferences of wild populations. The capacity of fast adaptation to changed environmental factors, facilitated by epigenetic mechanisms, could prove particularly useful in the face of rapid anthropogenic climate change. However, due to the high ecological and economic value of this kelp, the implication of changes in methylation due to cultivation, and especially the possibility of differential chloroplast methylation influencing photosynthetic capacity, need to be further assessed. Photosynthetic capacity is directly linked to primary production, and the capability to build up and maintain biomass. Processes impacting the chloroplast methylome would have implications for both cultivated and wild populations. In cultivars, the importance of matching hatchery and cultivation conditions to origin has been implied by our results. Furthermore, due to the significant differences observed in the methylomes of cultivated *S. latissima*, it might be possible to detect an epigenetic ‘fingerprint’ of cultivated specimen in the future, to detect escapees to the wild. Regarding the current trend in northward shift, the further north populations of *Saccharina latissima* settle, the more pressure the months of darkness during polar night will pose (Scheschonk et al. 2019). Photosynthetic pigments were shown to be functional throughout the prolonged darkness of the Arctic winter (Scheschonk et al 2019), indicating the chloroplasts of High Arctic specimen retained photosynthetic capacity during polar night. In which context differential methylation of chloroplast DNA facilitated by differences in the eco-phenotypes plays a role in this context will be of interest, especially regarding populations adapted to light conditions at more southern distributional habitats, shifting northwards. In *Saccharina latissima*, phenotypic variation among populations is often driven by phenotypic plasticity, and the species shows high plasticity in response to changing abiotic factors (Bolton & Lüning 1982; Diehl and Bischof 2021). So far, research on differences between origins in *S. latissima* has focused on genetic differentiation, with significant genetic variation between populations (Møller Nielsen et al. 2016; Paulino et al. 2016; Guzinski et al. 2020). Epigenetic differences have largely been unexplored as an alternative source of molecular variation between *S. latissima* populations. The differences we observed between origins within the laboratory and the field samples indicate that the eco-phenotype approach (Scheschonk et al 2021) is applicable to the organellar chloroplast genome as well. Furthermore, our data suggests that the influence of cultivation seems to be at least partially origin-specific, or that the influence of cultivation and origin on the methylome are interconnected. Epigenetic modifications have been found to be under some degree of genetic control, but can also be triggered by environmental factors such as temperature changes (Boyko et al. 2010; Correia et al. 2013). Hence, the role of genetic differences controlling the epigenetic responses reported here needs to be assessed in further detail in future studies.

## Data availability

Raw sequencing data of this study are available at NCBI SRA BioProject PRJNA809008.

## Acknowledgements

We thank S. Niedzwiedz (University of Bremen, Germany) for securing the field samples, and G. Hoarau and H. Reiss (Nord University, Bodø, Norway) for providing lab space and sequencer. Thank you, M. Kopp (Nord University, Bodø, Norway), for the support during the sequencing procedure, and for preparing some of the libraries. Thanks to I. Bartsch and A. Wagner (Alfred Wegener Institute for Marine and Polar Research, Bremerhaven, Germany) for providing and hatching the sporophyte cultures.

## Author Contributions

LS designed and conducted the experiment. AN, LS and AJ analysed and interpreted the data. LS and AN wrote the manuscript. All authors read and commented on previous versions of the manuscript, and approved the final version.

## Supplementary

### Suppl. Figures

**Supp. Fig. 1:**
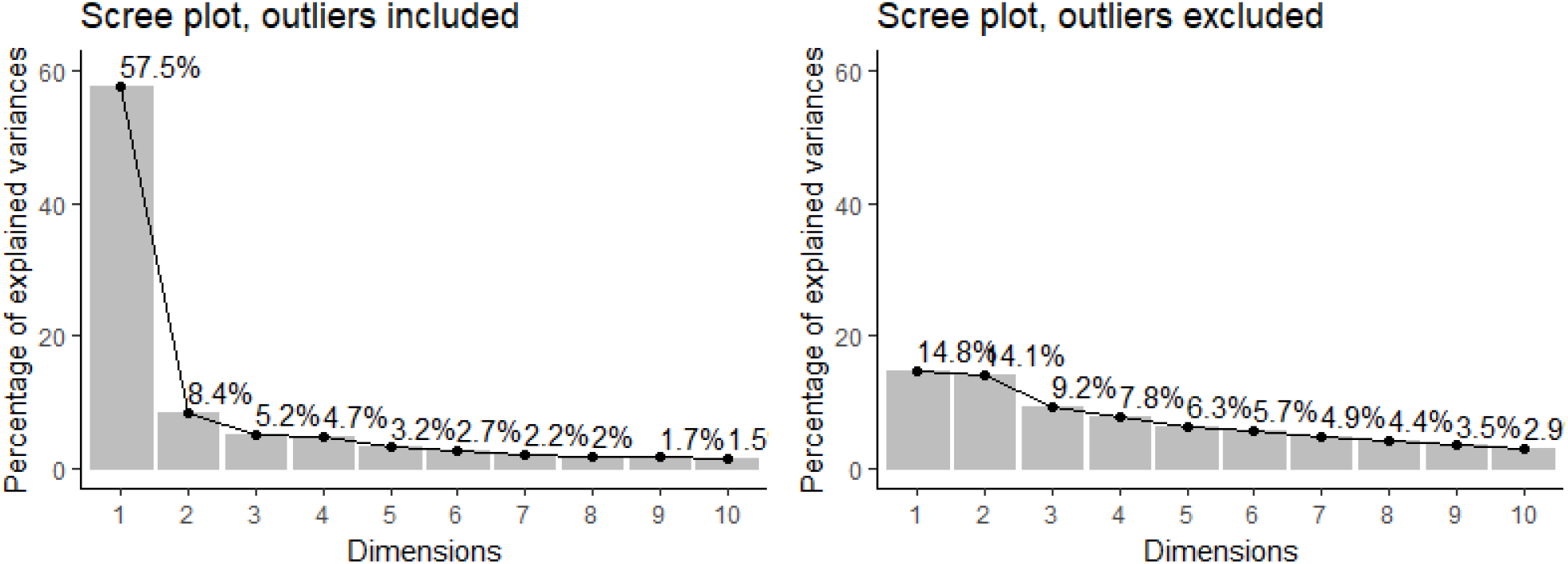
Scree plots showing the amount of variance explained by each dimension. On the left including all samples, on the right when removing two outliers.

**Supp. Fig. 2:**
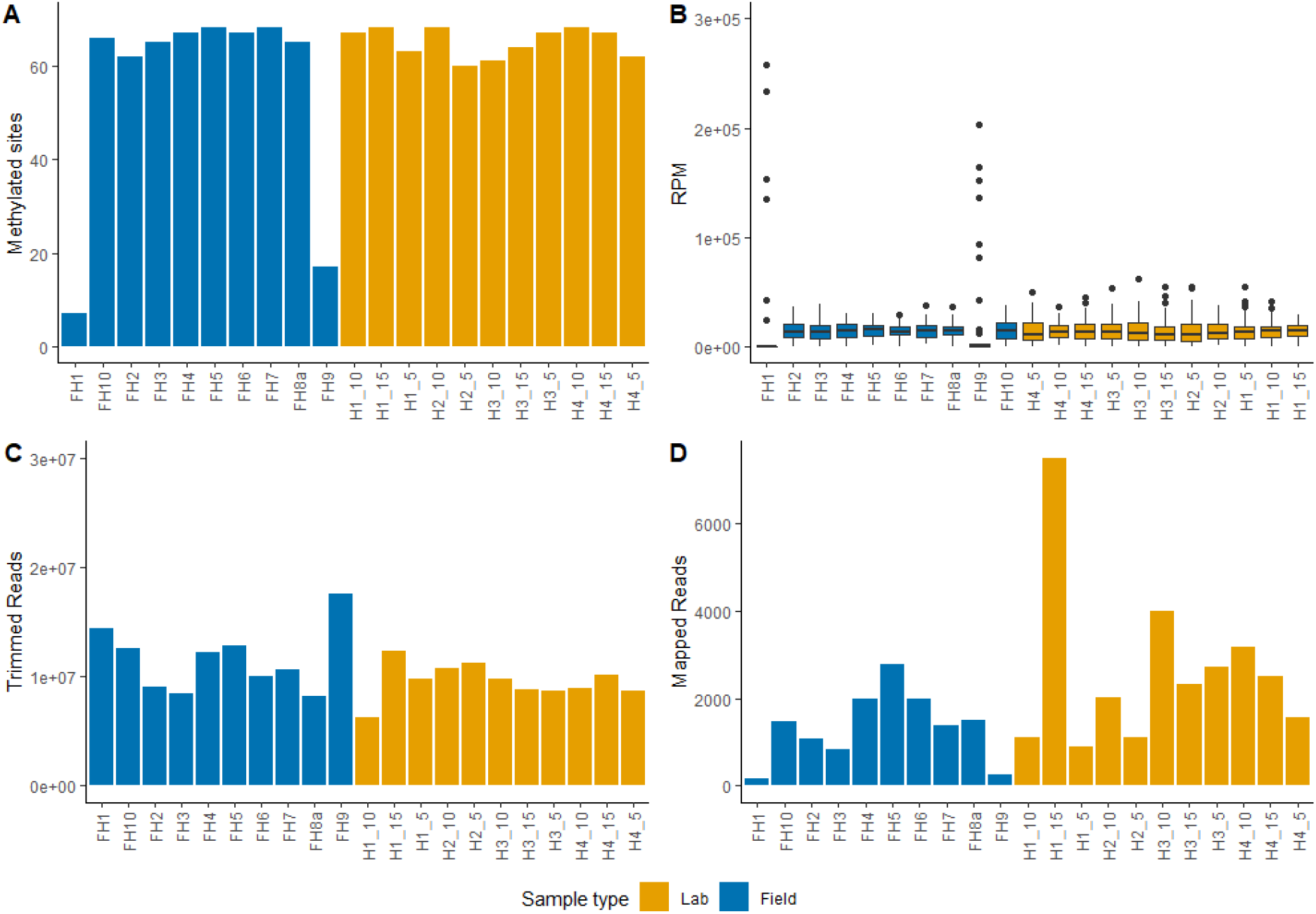
Sequencing output and methylation characteristics for each sample from Helgoland. Plots were used for outlier detection for PCA. A: Number of methylated sites (with more than two reads) for each sample. B: Boxplot showing reads per million (RPM) for each sample, with each point representing one methylated site. C: The number of sequenced and quality-trimmed reads for each sample. D: Numbers of quality-trimmed reads that mapped back to the chloroplast genome of *Saccharina latissima*.

**Supp. Fig. 3:**
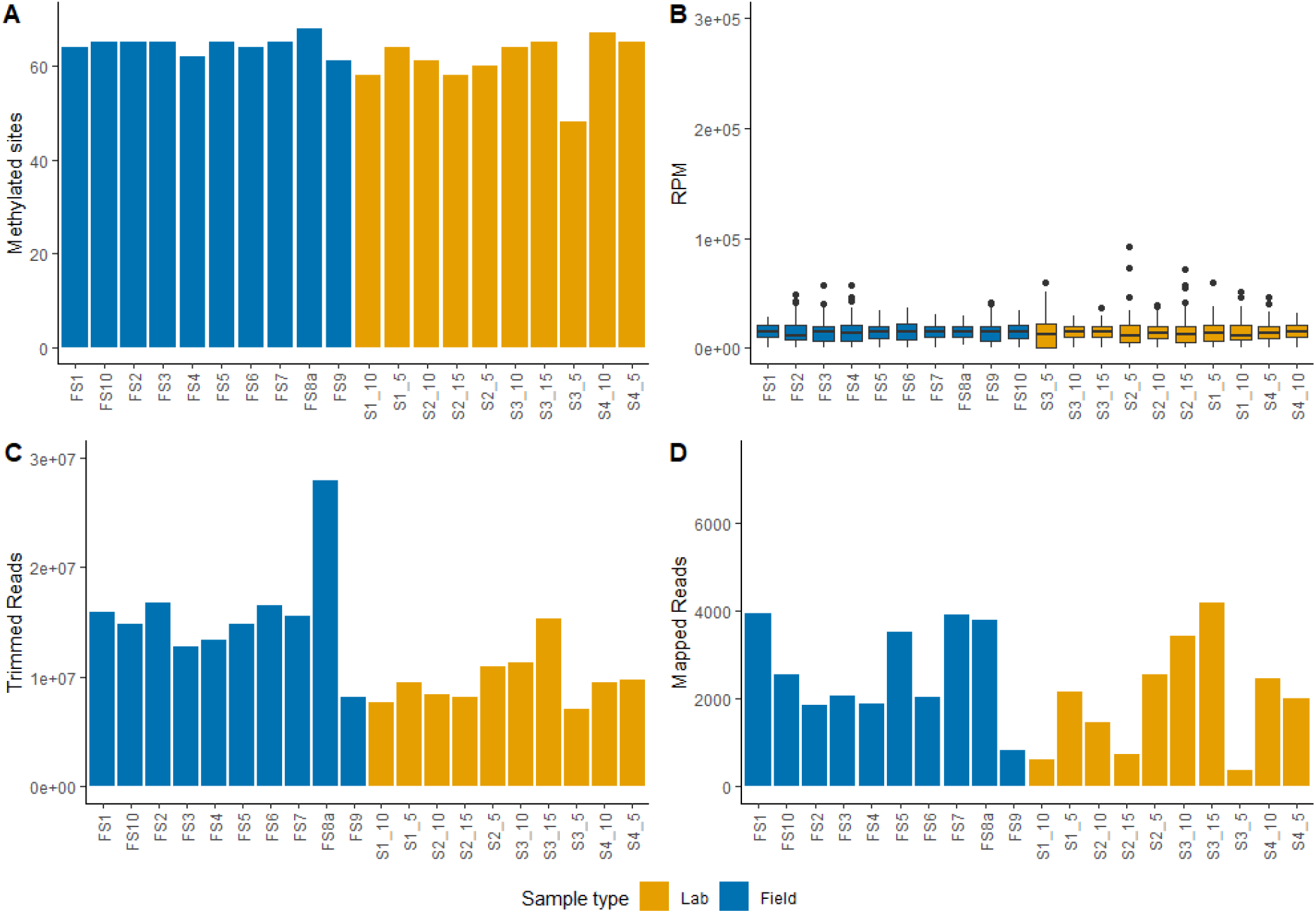
Sequencing output and methylation characteristics for each sample from Spitsbergen. Plots were used for outlier detection for PCA. A: Number of methylated sites (with more than two reads) for each sample. B: Boxplot showing reads per million (RPM) for each sample, with each point representing one methylated site. C: The number of sequenced and quality-trimmed reads for each sample. D: Numbers of quality-trimmed reads that mapped back to the chloroplast genome of *Saccharina latissima*.

**Supp. Fig. 4:**
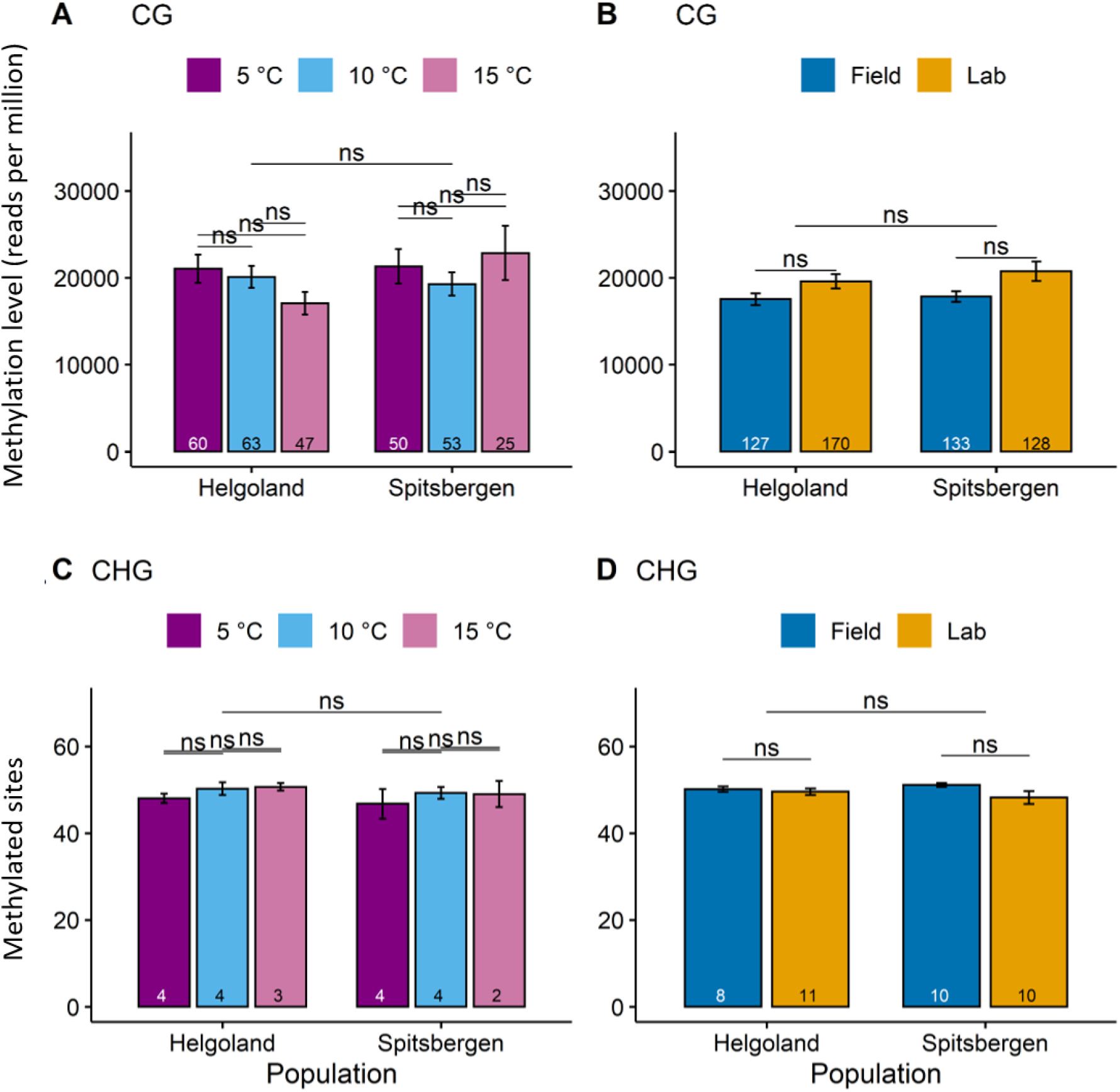
Differences in methylated sites and methylation level between conditions. Methylation levels as reads per million (RPM) for CHG sites (A, B), and methylated CG sites (A, B) between populations and temperatures in laboratory samples (A, C) and compared between lab and field samples (B, D). ns: not significant. Numbers at the bottom of the bars indicate sample size.

**Supp. Fig. 5:**
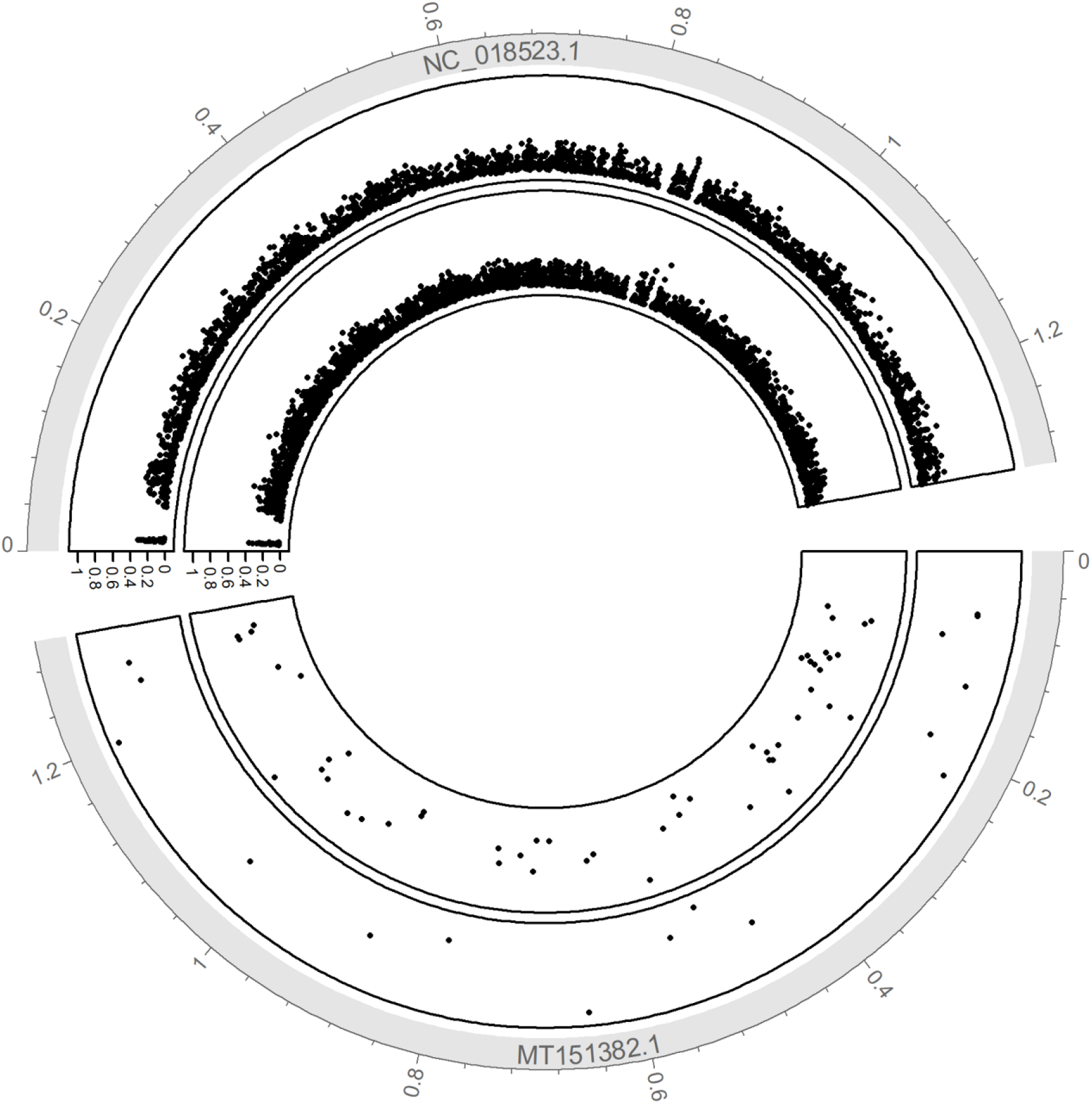
Overview of the chloroplast methylome in *S. latissima* (MT151382) and *S. Japonica* (NC_0198523.1). *S. japonica* data from Teng et al. 2021. Numbers on the outside of the circle represent locations within the genomes. Numbers on y-axes show the methylation levels at each site. The inner circle shows CHG methylation, and the outer circle shows CG methylation. Numbers along the outer circles refer to megabases along the chloroplast genomes.

### Suppl. Tables

**Suppl. Table 1:** Fertilisation scheme of the laboratory male (♂) and female (♀) gametophyte cultures from Helgoland (HG1-4) and Spitsbergen (SG1-4), resulting in the cultures (Helgoland (H1- 4) and Spitsbergen (S1-4) raised under laboratory conditions.

|  |  |  |  |  |  |  |  |
| --- | --- | --- | --- | --- | --- | --- | --- |
| Gametophyte | HG2 ♀ | HG3 ♀ | HG4 ♀ | SG1 ♀ | SG2 ♀ | SG3 ♀ | SG34 ♀ |
| Cultures | X | X | X | X | X | X | X |
|  | HG2 ♂ | HG3 ♂ | HG4 ♂ | SG1 ♂ | SG2 ♂ | SG3 ♂ | SG4 ♂ |
| Sporophyte | H2 | H3 | H4 | S1 | S2 | S3 | S4 |
| Culture |  |  |  |  |  |  |  |

**Suppl. Table 2.** Sites differentially methylated (p < 0.05) between Helgoland and Spitsbergen populations when including all samples in the comparison. Positive Log2FC values indicate increased methylation in Spitsbergen compared to Helgoland, while negative values indicate higher methylation in Helgoland samples.

| Location | Log 2FC | Log2F C std error | Adjusted p value | Gene ID | Gene name | Region | Product | Sequence context |
| --- | --- | --- | --- | --- | --- | --- | --- | --- |
| 21922-21953 | 1.04 | 0.196 | 2.66E-06 | QKE47428.1 | rpoB | gene | DNA-directed RNA polymerase beta chain | CHG |
| 68544-68575 | 0.933 | 0.302 | 0.0274 | QKE47473.1 | ycf3 | gene | photosystem I assembly protein | CHG |
| 39358-39389 | 0.456 | 0.159 | 0.0475 | QKE47440.1 | groEL | gene | 60 kDa chaperonin | CHG |
| 51728-51760 | -4.00 | 0.903 | 0.000164 | QKE47458.1 | psaA | promoter | photosystem I P700 apoprotein A1 | CG |
| 9045-9077 | -4.17 | 0.77 | 2.05E-06 | QKE47418.1 | atpA | gene | ATP synthase CF1 alpha chain | CG |
| 55344-55376 | -4.96 | 0.644 | 9.14E-13 | QKE47459.1 | psaB | gene | photosystem I P700 apoprotein | CG |
|  |  |  |  |  |  |  | A2 |  |

**Suppl. Table 3.** Sites with differential DNA methylation (p < 0.05) between Helgoland and Spitsbergen populations when comparing only field samples. Positive Log2FC values indicate increased methylation in Spitsbergen compared to Helgoland, while negative values indicate higher methylation in Helgoland samples.

| Location | Log2FC | Log2FC std error | Adjusted p value | Gene ID | Gene name | Region | Product | Sequence context |
| --- | --- | --- | --- | --- | --- | --- | --- | --- |
| 21922-21953 | 0.867 | 0.256 | 0.0117 | QKE47428.1 | rpoB | gene | DNA-directed RNA polymerase beta chain | CHG |
| 39358-39389 | 0.776 | 0.222 | 0.0104 | QKE47440.1 | groEL | gene | 60 kDa chaperonin | CHG |
| 75619-75650 | 0.727 | 0.206 | 0.0104 | QKE47479.1 | cbbX | gene | cfxQ-like protein | CHG |
| 55567-55598 | -0.528 | 0.173 | 0.0259 | QKE47459.1 | psaB | gene | photosystem I P700 apoprotein A2 | CHG |
| 51728-51760 | -3.15 | 0.994 | 0.0209 | QKE47458.1 | psaA | promoter | photosystem I P700 apoprotein A1 | CG |
| 55344-55376 | -4.49 | 1.21 | 0.0104 | QKE47459.1 | psaB | gene | photosystem I P700 apoprotein A2 | CG |

**Suppl. Table 4.** Sites with differential DNA methylation (p < 0.05) between samples originating from Helgoland or Spitsbergen when comparing only laboratory samples. Positive Log2FC values indicate increased methylation in Spitsbergen compared to Helgoland, while negative values indicate higher methylation in Helgoland samples.

| Location | Log2FC | Log2FC std error | Adjusted p value | Gene ID | Gene name | Region | Product | Sequence context |
| --- | --- | --- | --- | --- | --- | --- | --- | --- |
| 68544-68575 | 1.43 | 0.457 | 0.023 | QKE47473.1 | ycf3 | gene | photosystem I assembly protein | CHG |
| 21922-21953 | 1.18 | 0.285 | 0.000824 | QKE47428.1 | rpoB | gene | DNA-directed RNA polymerase beta chain | CHG |
| 9045-9077 | -5.24 | 0.701 | 2.71E-12 | QKE47418.1 | atpA | gene | ATP synthase CF1 alpha chain | CG |
| 51728-51760 | -5.24 | 1.43 | 0.00438 | QKE47458.1 | psaA | promoter | photosystem I P700 apoprotein A1 | CG |
| 55344-55376 | -5.68 | 0.639 | 4.27E-17 | QKE47459.1 | psaB | gene | photosystem I P700 apoprotein A2 | CG |

**Suppl. Table 5.** Sites differentially methylated (p < 0.05) between laboratory and field samples. Positive Log2FC values indicate higher methylation in laboratory samples compared to field samples.

| Location | Log2FC | Log2FC<br>std error | Adjusted<br>p value | Gene ID | Gene<br>name | Region | Product | Sequence<br>context |
| --- | --- | --- | --- | --- | --- | --- | --- | --- |
| 88361-88392 | 2.27 | 0.479 | 0.000137 | QKE47493.1 | rpl3 | gene | 50S ribosomal protein L3 | CHG |
| 15235-15266 | 1.16 | 0.282 | 0.000659 | QKE47426.1 | rpoC2 | gene | DNA-directed RNA polymerase beta' chain | CHG |
| 88373-88404 | 0.987 | 0.298 | 0.00741 | QKE47493.1 | rpl3 | gene | 50S ribosomal protein L3 | CHG |
| 17309-17340 | 0.952 | 0.31 | 0.0124 | QKE47426.1 | rpoC2 | gene | DNA-directed RNA polymerase beta' chain | CHG |
| 64927-64959 | 0.752 | 0.173 | 0.000294 | QKE47469.1 | petA | gene | cytochrome f | CG |
| 34231-34262 | 0.598 | 0.198 | 0.0134 | QKE47436.1 | secA | promoter | preprotein translocase subunit A | CHG |
| 102711-102742 | 0.591 | 0.172 | 0.00654 | QKE47520.1 | tufA | gene | translation elongation factor Tu | CHG |
| 124685-124717 | 0.561 | 0.181 | 0.0123 | QKE47544.1 | ycf39 | gene | NmrA domain-containing protein | CG |
| 46721-46753 | 0.551 | 0.195 | 0.0237 | QKE47451.1 | ycf42 | gene | putative peroxiredoxin ycf42 | CG |
| 32980-33011 | 0.514 | 0.143 | 0.00416 | QKE47436.1 | secA | gene | preprotein translocase subunit A | CHG |
| 102611-102643 | 0.49 | 0.175 | 0.0241 | QKE47520.1 | tufA | gene | translation elongation factor Tu | CG |
| 22542-22574 | 0.439 | 0.161 | 0.0264 | QKE47428.1 | rpoB | gene | DNA-directed RNA polymerase beta chain | CG |
| 75988-76019 | -0.435 | 0.158 | 0.0253 | QKE47479.1 | cbbX | gene | cfxQ-like protein | CHG |
| 104220-104251 | -0.567 | 0.173 | 0.00741 | QKE47523.1 | ftsH | gene | ATP-dependent zinc metalloprotease FtsH | CHG |
| 127333-127364 | -0.641 | 0.191 | 0.00741 | QKE47546.1 | ilvB | gene | acetolactate synthase large subunit | CHG |
| 95574-95605 | -0.728 | 0.16 | 0.000183 | QKE47509.1 | rps5 | gene | 30S ribosomal protein S5 | CHG |

**Suppl.Table 6:**
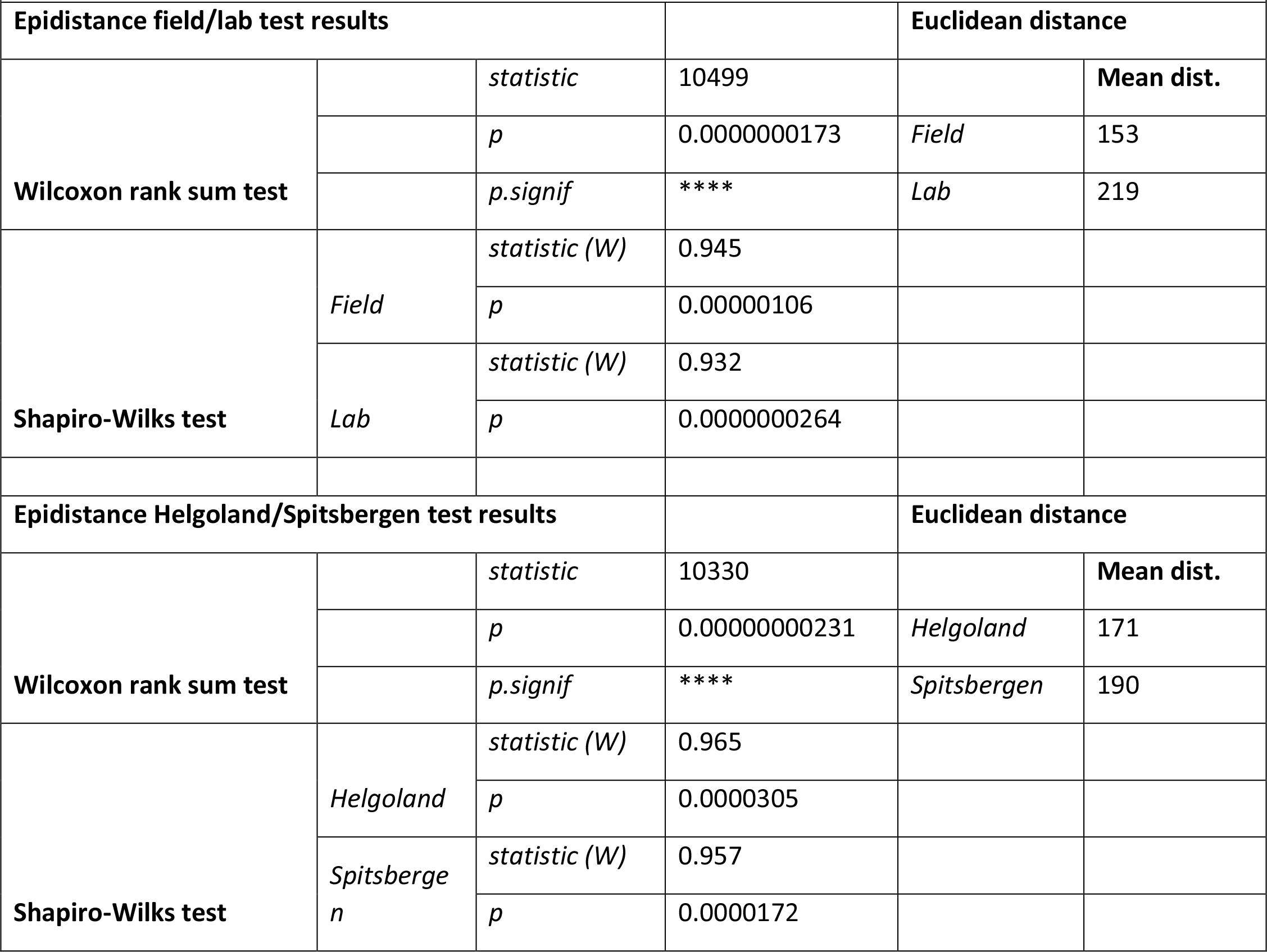
Statistics results of the epigenetic distance analysis.

| Suppl.Table 6: Statistics results of the epigenetic distance analysis. |  |  |  |  |  |
| --- | --- | --- | --- | --- | --- |
| Epidistance field/lab test results |  |  |  | Euclidean distance |  |
| Wilcoxon rank sum test |  | *statistic* | 10499 |  | Mean dist. |
|  |  | *p* | 0.0000000173 | *Field* | 153 |
|  |  | *p.signif* | **** | *Lab* | 219 |
| Shapiro-Wilks test | *Field* | *statistic (W)* | 0.945 |  |  |
|  |  | *p* | 0.00000106 |  |  |
|  | *Lab* | *statistic (W)* | 0.932 |  |  |
|  |  | *p* | 0.0000000264 |  |  |
| Epidistance Helgoland/Spitsbergen test results |  |  |  | Euclidean distance |  |
| Wilcoxon rank sum test |  | *statistic* | 10330 |  | Mean dist. |
|  |  | *p* | 0.00000000231 | *Helgoland* | 171 |
|  |  | *p.signif* | **** | *Spitsbergen* | 190 |
| Shapiro-Wilks test | *Helgoland* | *statistic (W)* | 0.965 |  |  |
|  |  | *p* | 0.0000305 |  |  |
|  | *Spitsberge*<br>*n* | *statistic (W)* | 0.957 |  |  |
|  |  | *p* | 0.0000172 |  |  |

Suppl.Table 7 see excel file ‘SupplTab7_8_9_Fig3_4_Tab2’ sheet ‘ST7 Origins’

Suppl.Table 8 see excel file ‘ SupplTab7_8_9_Fig3_4_Tab2’ sheet ‘ST8 Temperature’

Suppl.Table 9 see excel file ‘ SupplTab7_8_9_Fig3_4_Tab2’ sheet ‘ST9 Cultivation’

